# A Novel Mechanism of Cardiomyopathy: Toxic Peptides Dysregulate Calcium Transport

**DOI:** 10.64898/2026.04.24.719962

**Authors:** Taylor A. Phillips, Jacob D. Cunningham, Mary D. Hernando, Jaroslava Seflova, Laura A. Sherer, Seby Edassery, Jonathan A. Kirk, Howard S. Young, Seth L. Robia

## Abstract

A hallmark of dilated cardiomyopathy (DCM) is calcium mishandling, including reduced transport activity of the SERCA calcium pump in cardiac muscle cells. This has focused attention on SERCA as mechanism of disease and potential therapeutic target. Previously, diminished SERCA activity has been attributed to decreased protein expression, but recent studies suggest SERCA levels are unchanged in DCM. Thus, another mechanism must be responsible for the deficit. Since proteolysis is increased and proteosome function is impaired in DCM, we reasoned that accumulation of toxic protein fragments may contribute to SERCA dysfunction. In particular, previous studies showed diverse species of hydrophobic α-helices can inhibit SERCA, so we hypothesized that SERCA may become congested with transmembrane peptides that mimic endogenous regulatory partners. We purified cell membranes from non-failing and DCM human ventricles and subjected them to mass spectrometry to identify protein species upregulated in DCM. Select candidates were screened for binding and inhibition of SERCA. Several small membrane proteins and membrane protein fragments bound avidly to SERCA and significantly reduced cellular calcium stores. The data suggest a novel pathophysiological mechanism in which transmembrane protein debris obstructs SERCA function and regulation, contributing to cardiac muscle dysfunction in heart failure.

## Introduction

Dilated cardiomyopathy (DCM) is a leading cause of heart failure characterized by thinning of ventricle walls, weak contractions, and sluggish relaxation. DCM has many genetic causes, or it may be acquired due to cardiac inflammation, but up to 40% of DCM is idiopathic. This has prompted intensive investigation of pathophysiological mechanisms contributing to DCM. At the cellular level, a key hallmark of DCM is disordered Ca handling^1-3^. Decreased Ca transport by the sarcoplasmic reticulum calcium ATPase (SERCA) results in decreased intracellular Ca stores, which contributes to poor contractility. Diminished Ca transport also causes slower muscle relaxation during diastole^3,4^, so both phases of the cardiac cycle are impacted by compromised SERCA function^3,5^. The decrease in SERCA function in DCM and other forms of heart failure has been attributed to a decrease in SERCA protein expression^6,7^, but other recent studies suggest that SERCA is still highly expressed^8^, so impaired cardiac Ca transport must be due to another undiscovered pathophysiological mechanism. This suspicion is supported by the failure of clinical trials that attempted to supplement transport function by adeno-associated virus gene delivery of SERCA2a to the myocardium^7^. We hypothesize that the fundamental problem of DCM is dysregulation of SERCA, so increasing SERCA expression into an environment of disordered cellular Ca regulatory mechanisms cannot address the transport deficiency.

In the healthy heart, the rate of Ca uptake into the sarcoplasmic reticulum (SR) is tightly regulated and responsive to neuroendocrine signaling. β-adrenergic stimulation enhances cardiac contractility through PKA-mediated phosphorylation of multiple targets including the archetypal regulator of SERCA, the microprotein phospholamban (PLB)^9,10^. Phosphorylation of PLB relieves its inhibition of SERCA, yielding faster Ca^2+^ reuptake and cardiac relaxation^10,11^.Notably, this response to β-adrenergic stimulation is decreased or absent in DCM, leading to a less dynamic range of cardiac contractility and worsening of the disease^12-14^. In addition to PLB, other small transmembrane peptides can interact with SERCA and regulate pump function. The members of this family of microproteins, or “regulins”, have a strong inhibitory effect on SERCA, with the exception of DWORF^15^, which activates SERCA directly^16-18^. DWORF also competitively displaces inhibitory PLB from the binding site on SERCA^19,20^. The regulins have relatively low sequence homology but they all have a hydrophobic, α-helical transmembrane domain that interacts with a cleft in the transmembrane region of the SERCA pump. SERCA’s hydrophobic binding cleft is broadly specific to accommodate the diverse regulins. Indeed, the wide range of possible binding partners was demonstrated by a study of the inhibitory potential of synthetic peptides, which showed simple sequences of leucine-alanine repeats could inhibit SERCA as effectively as native regulatory partners^21,22^. In view of the broad selectivity of the regulatory site, we hypothesized that SERCA might be dysregulated by improper protein-protein interactions in the chaotic proteome of the failing heart.

In DCM, proteolysis is greatly increased, overwhelming the proteosome and leading to partially degraded proteins^23-26^. Calpains, caspases and other proteases are expressed, many of which are localized to the SR membrane^27-31^. SR proteins are partially protected by the lipid bilayer of the SR membrane, but luminal and cytosolic aspects are exposed, so proteolysis generates persistent transmembrane fragments. Such hydrophobic, single pass transmembrane peptides possess the structural determinants to bind the SERCA pump. In addition to changes in protein degradation, the synthesis of many proteins changes in DCM^32,33^. We hypothesize that increased proteolysis and altered protein expression together lead to the production of SERCA-inhibitory peptides that dysregulate cellular Ca handling in heart failure.

## Results

### Upregulation of Small Peptides in Human DCM Samples

To determine how disruption of the proteome in the failing heart may alter the membrane protein millieu and impact SERCA regulation, we obtained specimens of human myocardium of explanted DCM hearts and compared them to non-failing human myocardium specimens. DCM human heart samples were obtained from the explanted hearts of patients receiving donor hearts. Echocardiograms were performed on all patients to validate the dilated phenotype, and hypertrophic and ischemic specimens were ruled out for this study. Non-failing samples were obtained from rejected donor hearts due to non-cardiac related reasons. **Fig. 1A** depicts the analytical workflow from proteomics to microscopy to activity measurements, successively reducing species to obtain highest likelihood candidates. To focus on transmembrane peptide species with high likelihood to interact with SERCA and modulate its function, we isolated SR membrane fractions using differential centrifugation of homogenized myocardium (**Fig. 1A**). Triplicate samples from each specimen were run on an SDS-PAGE gel to restrict to proteins of size less than 25 kD (**Fig. 1B**, dotted line). This size cut-off is appropriate to select single-span transmembrane species compatible with the micropeptide regulatory binding cleft on SERCA. Gel pieces from each sample lane were subjected to in-gel trypsin digestion. Digested peptides were analyzed by high performance liquid chromatography (HPLC) and mass spectrometry to compare the relative abundance of SR membrane proteins in DCM and NF samples. Over 1800 proteins were detected across all sample groups. 92 of these were significantly upregulated in DCM groups compared to NF controls (**Fig. 1C**).

**Figure 1.**
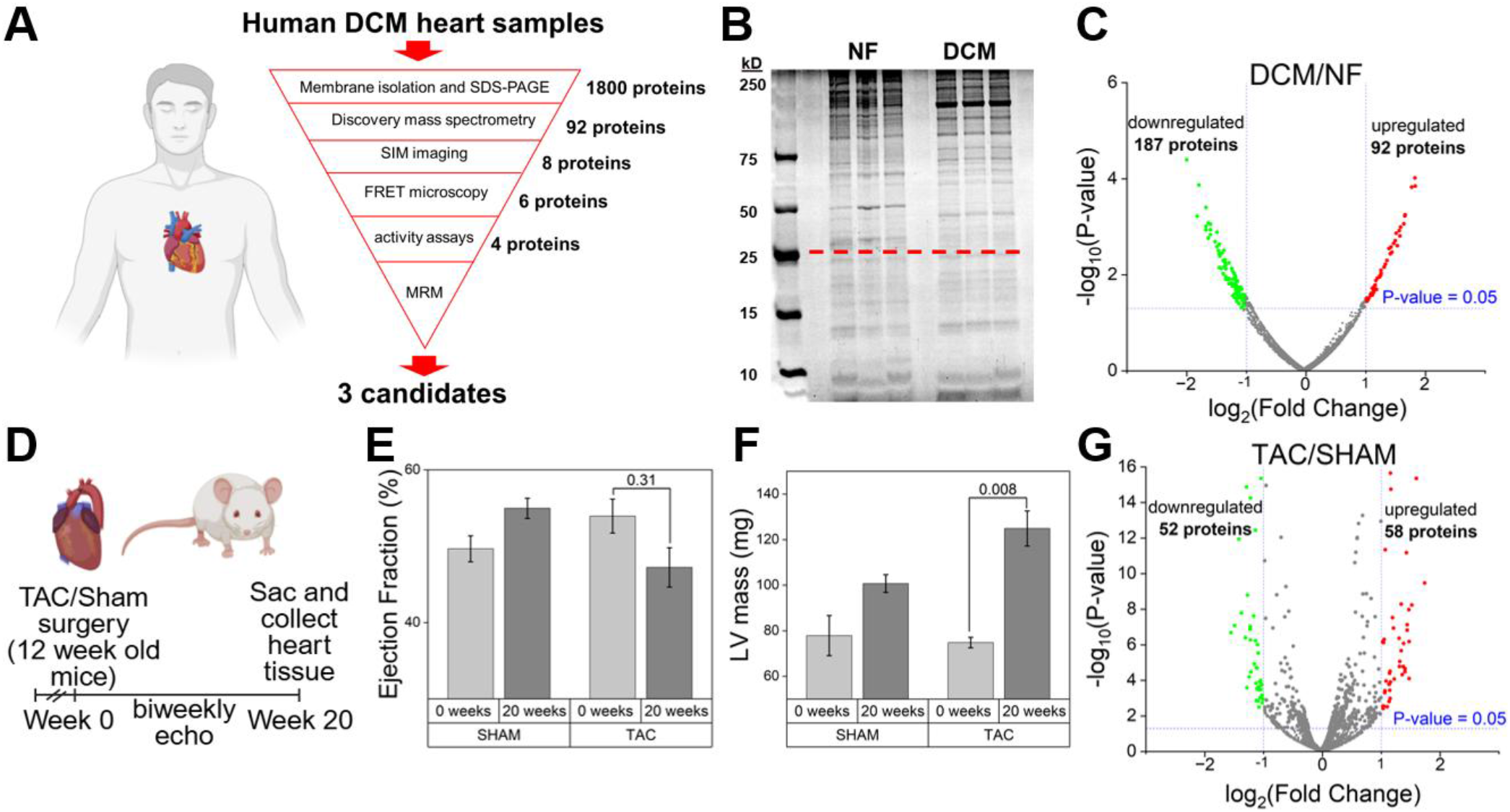
**A)** Experimental design showing process of identifying SERCA-inhibitory candidates in human DCM samples compared to non-failing control heart samples. **B)** SDS-PAGE gel showing total protein expression of the membrane fraction of non-failing and DCM human heart samples. Gel was cut below 25 kD marker. **C)** Volcano plot of differentially expressed proteins detected via mass spectrometry. Red dots represent over 3-fold upregulated in DCM and green dots over 3-fold downregulated in DCM compared to non-failing control. **D)** Experimental design of TAC and sham surgeries performed on 12-week-old mice followed by bi-weekly echocardiograms and collection of cardiac tissue. E) No change in ejection fraction (%) in TAC or sham animals 20 weeks post surgery. N=3 for each group. F) Increased left ventricle mass (mg) 20 weeks post TAC surgery. No change in SHAM mice. N=3 for each group. **G)** Volcano plot of differentially expressed proteins detected via mass spectrometry. Red dots represent over 3-fold upregulated in TAC mouse heart samples and green dots over 3-fold downregulated in TAC samples compared to sham control.

MS detected many peptides derived from proteins larger than the 25 kD cut-off, consistent with proteolysis of larger proteins into smaller species with greater electrophoretic mobility. Other peptides identified by MS were from proteins with a native size of 25 kD or less, so we presume the increase in those proteins was due to increased transcription/translation. Peptide sequences that included transmembrane spans or were immediately adjacent to transmembrane regions in the protein primary sequence were considered candidates that might bind SERCA and modulate its function. For large, multi-pass membrane proteins that were detected among low molecular weight species on the gel, we focused on the transmembrane region or regions nearest the peptide that was detected by MS. For smaller, single-pass proteins, the entire protein was regarded as a possible SERCA-binding species. To refine the list of candidates, we examined primary sequences for homology with the diverse micropeptides known to regulate SERCA. 8 candidate sequences of various lengths were selected for further screening and were fused to the C-terminus of YFP for localization and FRET studies (**Table 1**). For candidates SLMAP, TMEM14C, TMEM50A, EMC6, VAMP3, and VAMP8, we selected transmembrane regions adjacent to the identified peptide sequence (**bolded on Table 1**). We used the entire sequence of two small, single span transmembrane proteins, SEC61β and SMIM13.

**Table 1.**
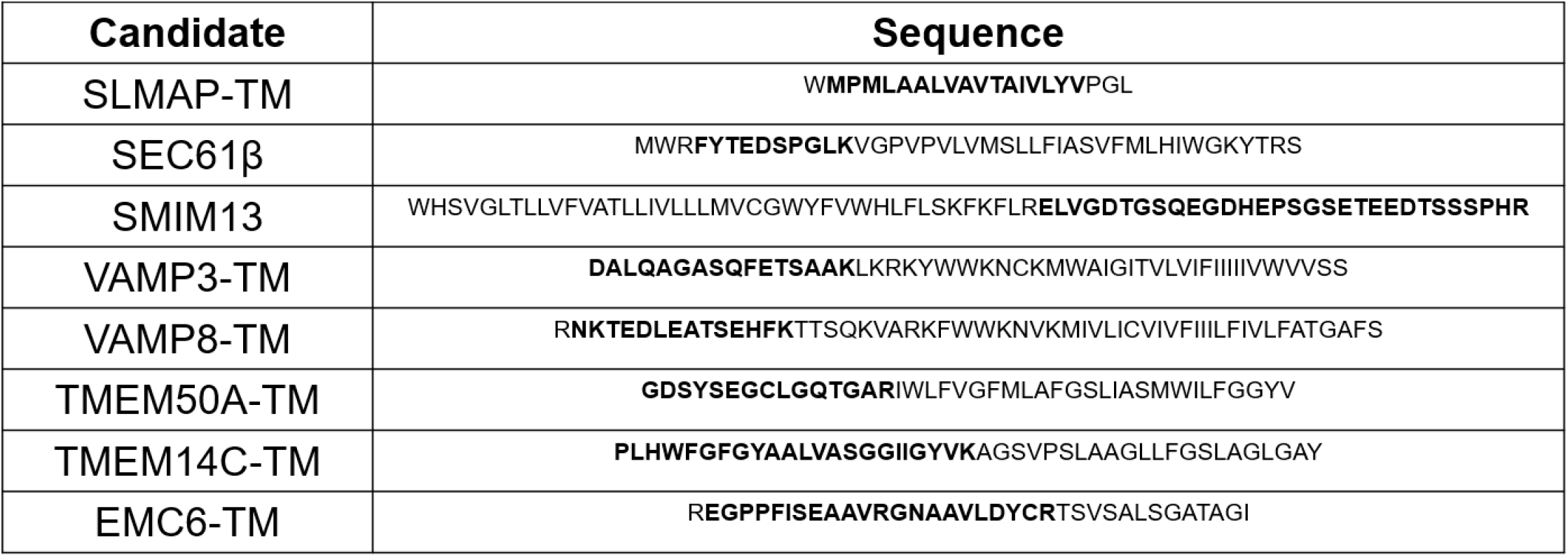
List of candidate peptides selected. Amino acid sequence showing transmembrane region of each protein and peptide fragment detected on mass spectrometry shown in bold.

A limitation of our analysis of explanted failing human hearts is that the specimens represent end-stage disease, making it difficult to disentangle initial pathogenic causes from later downstream consequences. To determine if the proteome changes we observed in late-stage DCM first occur at an earlier stage of disease, we mimicked mild heart failure with a modestly restrictive transaortic constriction (TAC) mouse model (**Fig. 1D**). TAC or sham surgeries were performed on 12-week-old wild type mice. Following surgery, bi-weekly echocardiograms were performed on mice until left ventricle remodeling and dilation was observed. 20 weeks post TAC surgery, there was no significant change in ejection fraction (**Fig. 1E**), however we observed a significant increase left ventricle mass signifying a mild hypertrophic phenotype (**Fig. 1F**). Animals were sacrificed and heart tissue was collected for fractionation of membrane and PAGE protein size restriction according to the same protocol used for human tissue specimens. MS results show many proteins were differentially expressed between TAC and sham groups (**Fig. 1G**), with significant overlap of candidate toxic species between the mouse model and the human DCM samples. Specifically, we observed SLMAP, EMC6, and many SMIMs/TMEMs were increased by 2-3 fold in TAC samples. The data suggest that candidate peptides are produced at an early stage of heart failure. Thus, they are not merely a feature of advanced disease and can be evaluated for possible contribution to the pathogenesis of DCM.

### Localization of Candidate Toxic Peptides

We reasoned that putative toxic peptides must have SR membrane localization to bind SERCA. We performed structured illumination microscopy (SIM) to determine the localization of candidates heterologously expressed as YFP-fusion proteins in HEK cells. We compared candidates to several ER-localized controls: SERCA and two of its native regulatory micropeptides, PLB and ALN (**Fig. 2A-C**). Of the candidates tested, 6 demonstrated ER localization (**Fig. 2D-I**) and co-localization with mCer-SERCA. Interestingly, VAMP3 and VAMP8 (**Fig. 2E-F**) demonstrated plasma membrane localization in addition to ER localization. We have previously observed a fraction of PLB expressed in the plasma membrane^34^; it is unclear whether this is a result of heterologous overexpression or represents a true functional role of PLB in the PM^35-37^. TMEM14C (**Fig. 2J**) demonstrated cytoplasmic localization and was not considered further as we reasoned that a candidate must co-localize with SERCA to regulate its activity.

**Figure 2.**
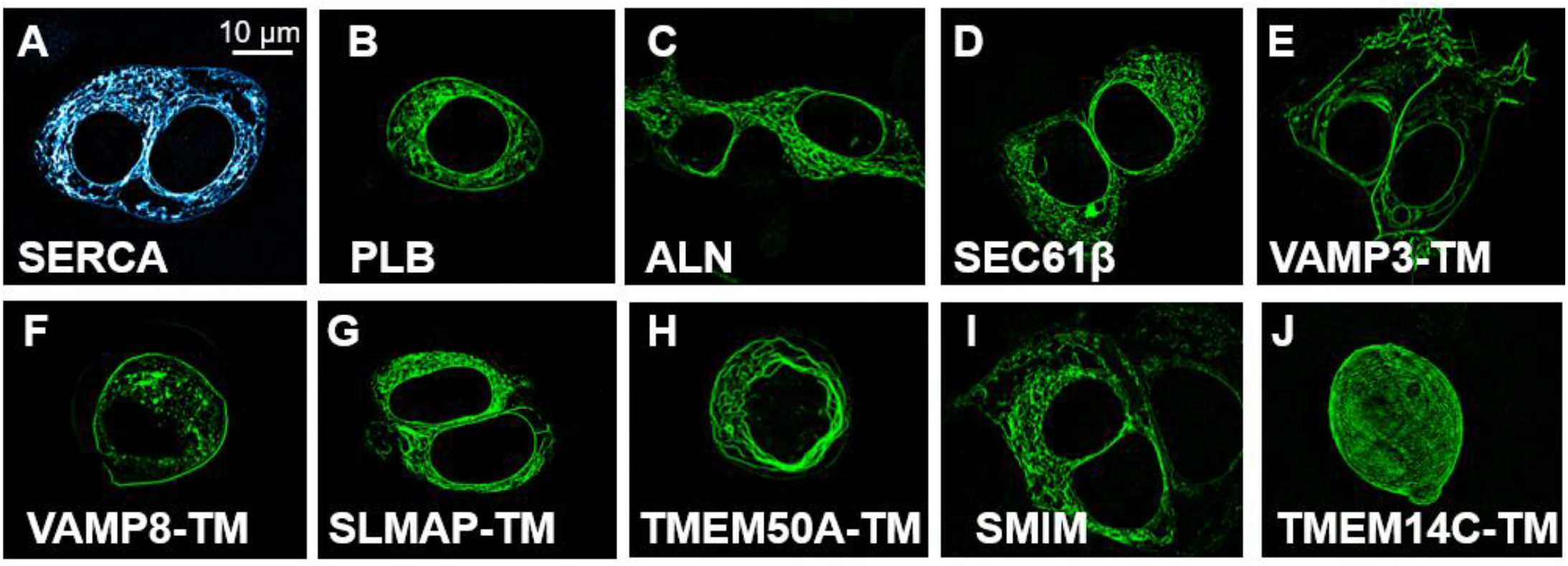
Structured illumination microscopy (SIM) images of transfected HEK 293T cells. SERCA, PLB, and ALN used as a positive control for ER localization. **A)** mCer tagged SERCA2a **B)** YFP tagged PLB **C)** YFP tagged ALN. **D)** SEC61 translocon subunit β, whole protein. **E)** YFP tagged vesicle-associated membrane protein 3, whole protein. **F)** YFP tagged vesicle-associated membrane protein 8, whole protein. **G)** YFP tagged sarcolemma associated protein, transmembrane (helix #1). **H)** YFP tagged transmembrane protein 50A, transmembrane [helix #4]. **I)** YFP tagged small integral membrane protein 13, whole protein **J)** YFP tagged transmembrane protein 14C, transmembrane (helix #2). Demonstrates ubiquitous expression.

### Quantifying binding affinity to SERCA

The observed increase in peptides with the structural features appropriate to bind SERCA was compatible with the hypothesis of toxic interactions with the pump. To test if the ER-localized candidates bind to SERCA in the intact membrane environment of living cells, FRET microscopy was performed as previously described^38,39^. Briefly, donor (mCer) and acceptor (YFP) tags were attached to SERCA and candidate peptides, respectively. The fluorophores were excited in donor, acceptor and FRET channels. Fluorescence intensity in each channel was quantified for every cell to calculate FRET efficiency. The FRET efficiency of each cell was plotted as a function of YFP fluorescence, taken as an index of acceptor protein expression. This relationship was well-described by a hyperbola of the form FRET% = (FRET_max_)([acceptor])/(K_D_+[acceptor]), where FRET_max_ is the maximum FRET efficiency at high protein expression levels (when all donors participate in FRET) and K_D_ is the relative dissociation constant, which is inversely related to the affinity of each candidate for SERCA. The K_D_ measured in this way is a relative value with arbitrary units (A.U.), so PLB and ALN, two known SERCA binding proteins, were compared to the candidate peptides. The SERCA-PLB interaction had a K_D_ of 1.8 A.U. (**Fig. 3B**) and SERCA-ALN had a K_D_ of 6 A.U. (**Fig. 3C**). Candidates yielding significantly higher values of K_D_ were considered to have poor SERCA-binding affinity, making them unlikely candidates to modulate SERCA function (particularly given presumed competition from endogenous regulatory micropeptides). To exclude poor-binding candidates with low affinity for SERCA, we eliminated from consideration those candidates with K_D_ above a cut-off criterion of 20 A.U. Four candidates (SEC61β, VAMP3, VAMP8 and SLMAP-TM) demonstrated acceptable binding affinity to SERCA (**Fig. 3D-G**), while candidates with poor binding affinities for SERCA, TMEM50A and SMIM (**Fig. 3H-I**), were excluded from further evaluation.

**Figure 3.**
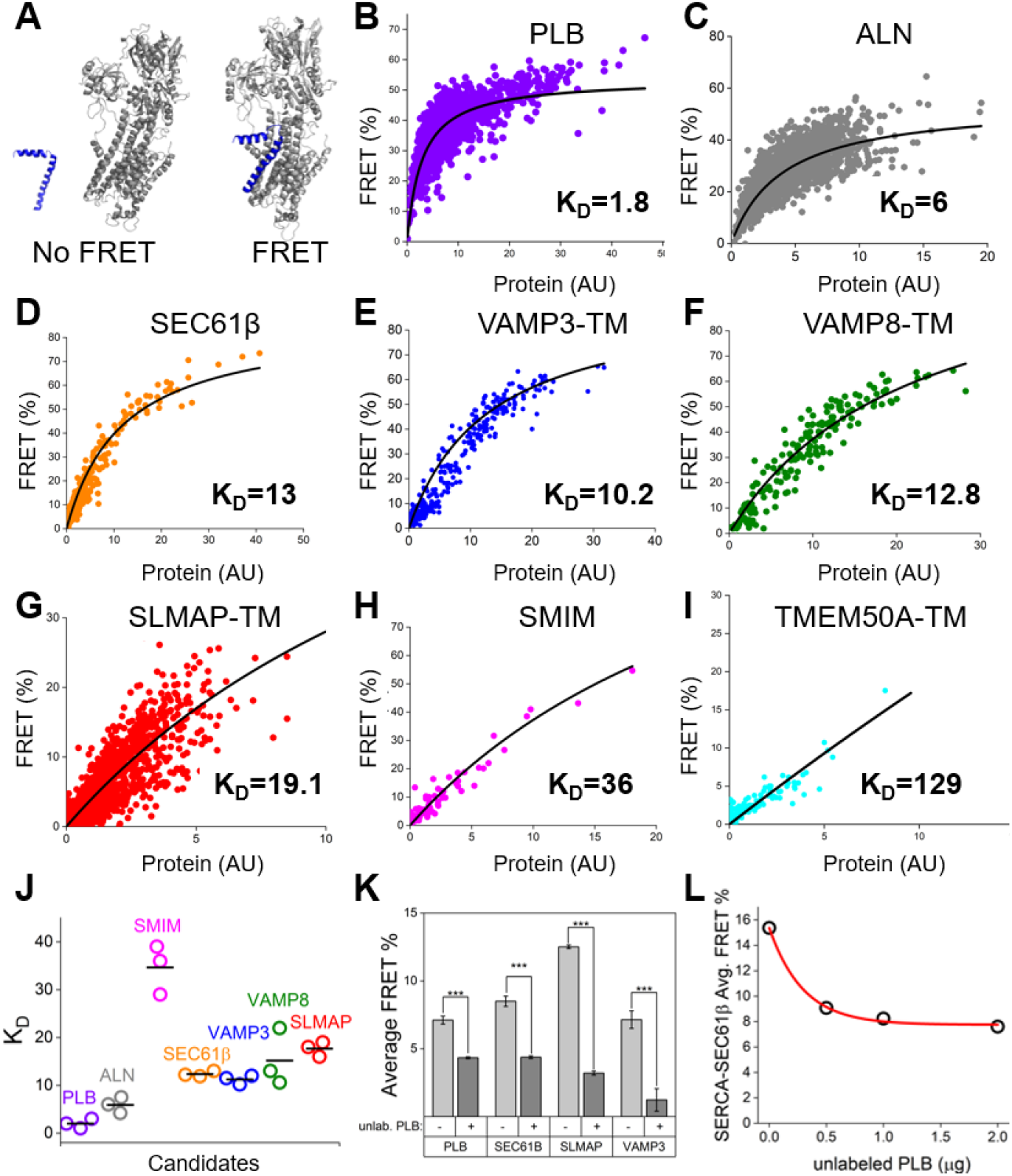
**A)** Binding of YFP-labeled PLB (blue) to Cer-labeled SERCA (grey) results in FRET. **B-I)** Binding curves of known SERCA inhibitors and selected poison peptide candidates. FRET (%) is quantified between mCer tagged SERCA and YFP tagged candidate with each point representing a different cell. Hyperbolic curve of best fit shown in black and K_D_, or relative binding affinity, displayed in red. **J)** Summary of K_D_ values for positive controls and candidates. **K)** Competition experiments with average FRET% between 0.25 μg SERCA and 0.5 μg candidate shown in red and with the addition of 10 μg unlabeled PLB shown (10x competitor) in blue. **L)** Average FRET between SERCA and SEC61β with increasing dosage of unlabeled PLB competitor.

Low K_D_ values are suggestive of an avid physical interaction between candidate peptides and SERCA, but we also endeavored to test the specificity of the observed candidate-SERCA interactions and explore the possibility of direct competition between candidates and PLB. We tested whether coexpression of unlabeled PLB reduced FRET from Cer-SERCA to YFP-labeled candidate peptides. We observed significant decreases in FRET efficiency after coexpression of PLB, suggesting peptide candidates compete directly with native micropeptides for the same regulatory binding site on SERCA (**Fig. 3J**). Competitive binding of unlabeled PLB also provides a way to estimate the contribution of non-specific FRET between donor-labeled SERCA and acceptor-labeled candidates that are concentrated near one another in the membrane, but not part of the same molecular complex. SERCA-candidate FRET decreased with increasing concentrations of PLB (**Fig. 3L**) to a minimum of 4-8%, which is similar to values we have measured in analogous competition experiments in previous studies^38^. According to the relationship described by Fung & Stryer^40^, this would correspond to a density of one acceptor-labeled protein per 1500 to 3500 lipids. For comparison, native sarcoplasmic reticulum protein levels are more concentrated than this, on the order of 1 protein per 750 lipids^41,42^. Thus, we do not consider the expression levels achieved in HEK cell to constitute “overexpression”, and we conclude that the interaction of SERCA with candidate poison peptides occurs at protein expression levels that are comparable to the native concentration of proteins in the SR membrane.

### The effect of candidate peptides on SERCA activity

Having established that candidate peptides interact with SERCA, we determined whether the peptide candidates were able to modulate SERCA activity. As an index of SERCA activity, we quantified ER calcium ([Ca^2+^]_ER_) uptake in live cells as previously described^43^. Briefly, HEK293T cells were transfected with SERCA, ryanodine receptor, and candidate peptides or known micropeptide regulators, together with GFP-rCepia1*er*, a ratiometric [Ca^2+^]_ER_ indicator (**Fig. 4A**). The fluorescence from rCepia1*er*, which increases at higher [Ca^2+^]_ER_, was normalized to GFP fluorescence and calibrated to quantify absolute [Ca^2+^]_ER_ (**Fig. 4B**). With no inhibitory peptides expressed, the [Ca^2+^]_ER_ was 350 µM. When PLB was co-expressed, [Ca^2+^]_ER_ decreased to 250 µM, indicative of decreased SERCA activity as a result of inhibition by PLB. Consistent with the known broad selectivity of SERCA’s regulatory site^22^, all candidates tested here displayed some inhibitory potency when co-expressed with SERCA (**Fig. 4C**). VAMP3 and VAMP8 decreased [Ca^2+^]_ER_ to 250 µM. SEC61β and SLMAP demonstrated the strongest inhibition of SERCA, lowering [Ca^2+^]_ER_ to below 200 µM. We observed a protein expression-dependent decrease in [Ca^2+^]_ER_ for control micropeptide PLB (**Fig. 4D**) and candidate poison peptides such as SEC61β (**Fig. 4E**) consistent with a saturable regulatory interaction with SERCA.

**Figure 4.**
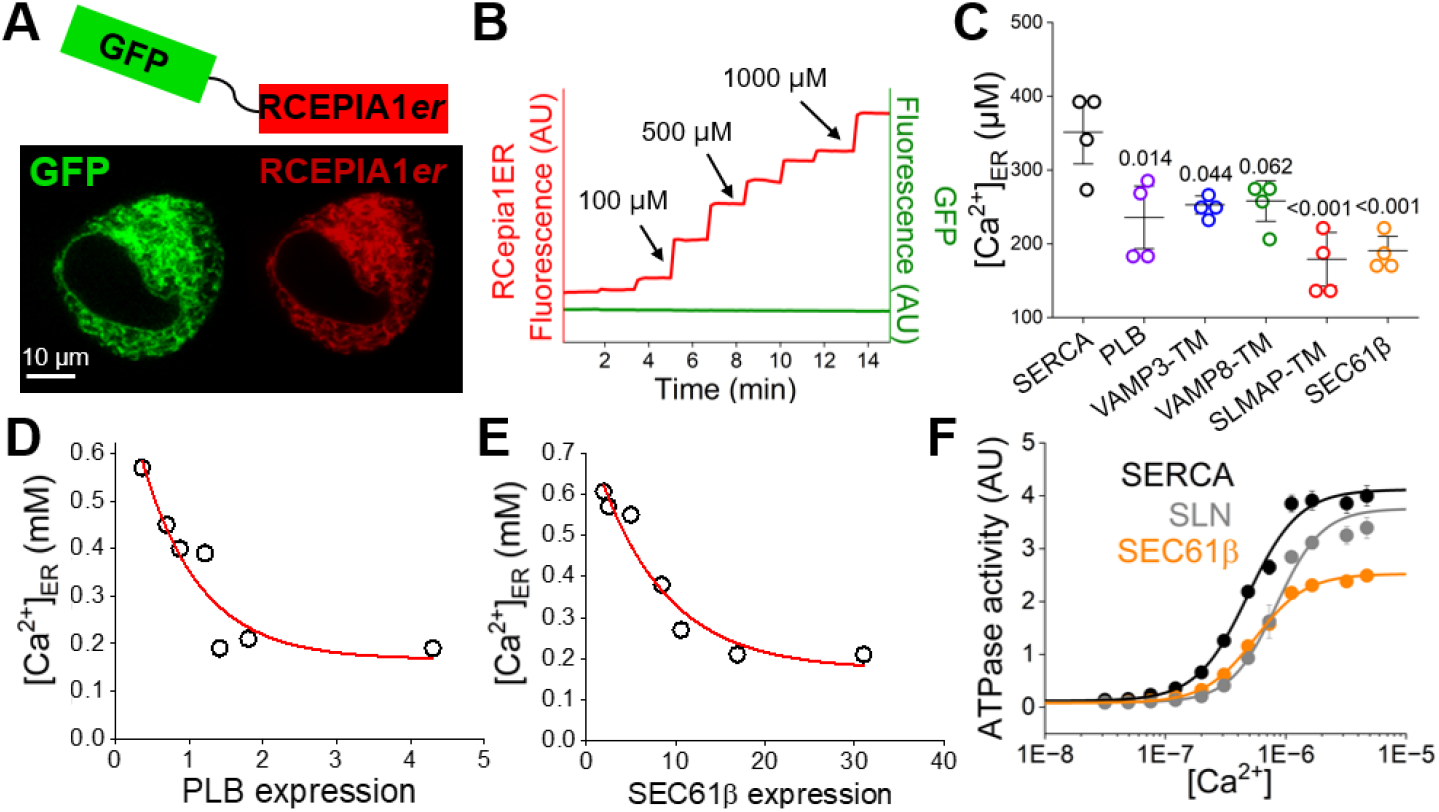
**A)** GFP-RCEPIA1*er*, used as ratiometric ER calcium indicator, consists of GFP fluorophore fused to RCEPIA1*er* with 21 amino acid linker between. Both fluorophores demonstrate ER localization in HEK293T cells. **B)** *RCEPIA1er* fluorescence increases as ER calcium concentration increase. GFP fluorescence remains constant, so can be used to measure of probe transfection. **C)** HEK293T cells transfected with SERCA, ryanodine receptor, peptide candidates and GFP-rCepia1*er*. Candidates compared to a SERCA negative control and PLB positive control. N=4 independent transfections with around 10,000 cells imaged per well. **D)** ER calcium concentration at increasing PLB expression levels, as determined by fluorescence intensity of transfected cells. **E)** ER calcium concentration at increasing SEC61β expression levels, as determined by fluorescence intensity of transfected cells. **F)** ATPase activity at various calcium concentrations for SERCA alone (black), SEC61β (orange), and SLN control (gray).

To determine whether candidate micropeptides altered cell calcium uptake by altering SERCA ATPase activity, SERCA and candidate peptides were reconstituted into proteoliposomes to measure the rate of ATP hydrolysis by the enzyme-coupled reaction assay based on optical quantification of production of NAD^+^ from NADH^45^. ATP hydrolysis was measured at varying calcium concentrations to quantify the K_Ca_, an indicator of SERCA calcium sensitivity, and V_max_, the maximal rate of hydrolysis. Compared to SERCA alone, SEC61β significantly reduced the V_max_ (**fig. 4F**) indicating slower enzymatic cycling SERCA at high Ca^2+^. The observed inhibition of V_max_ was even more pronounced than that observed for SLN. The other two tested candidates, VAMP3 and SLMAP, displayed an apparent decrease in the mean V_max_, though this difference did not achieve statistical significance. Overall, we conclude that several candidate poison peptides reduce Ca^2+^ uptake by SERCA, and we find SEC61β inhibits SERCA by reducing the pump’s maximum cycling rate.

### Quantification of candidate abundance in DCM

Discovery MS experiments revealed the upregulation of all screened peptides in DCM, but did not identify changes to full length protein from which peptide fragments are produced. To investigate changes to full length proteins and their native binding partners, we used western blotting (WB). WB demonstrates a decrease in expression of full-length SLMAP in DCM (**Fig. 5A**). The antibody used was not sensitive to TM fragment so only the bands at 95 kD were analyzed for relative abundance change (**Fig. 5E**). This decrease in full-length SLMAP, and increase of small peptide fragment from discovery MS, suggests degradation of the protein in DCM. SEC61β, a small transmembrane peptide, showed an increased expression in DCM (**Fig. 5B**) quantified to have around a 1.5x increase compared to NF controls (**Fig. 5F**). Interestingly, WB revealed that the expression of the large pore-forming subunit SEC61α was decreased in DCM compared to NF (**Fig. 5C and G**). The β subunit is typically associated with this larger α complex but in DCM the stoichiometry of the complex shifts which leaves more unbound β subunit to interact with SERCA. Western blots of full-length VAMP 3 showed significant increase in expression in DCM (**Fig. 5D and H**). This suggests that upregulation of this candidate may be due to transcriptional upregulation.

**Figure 5.**
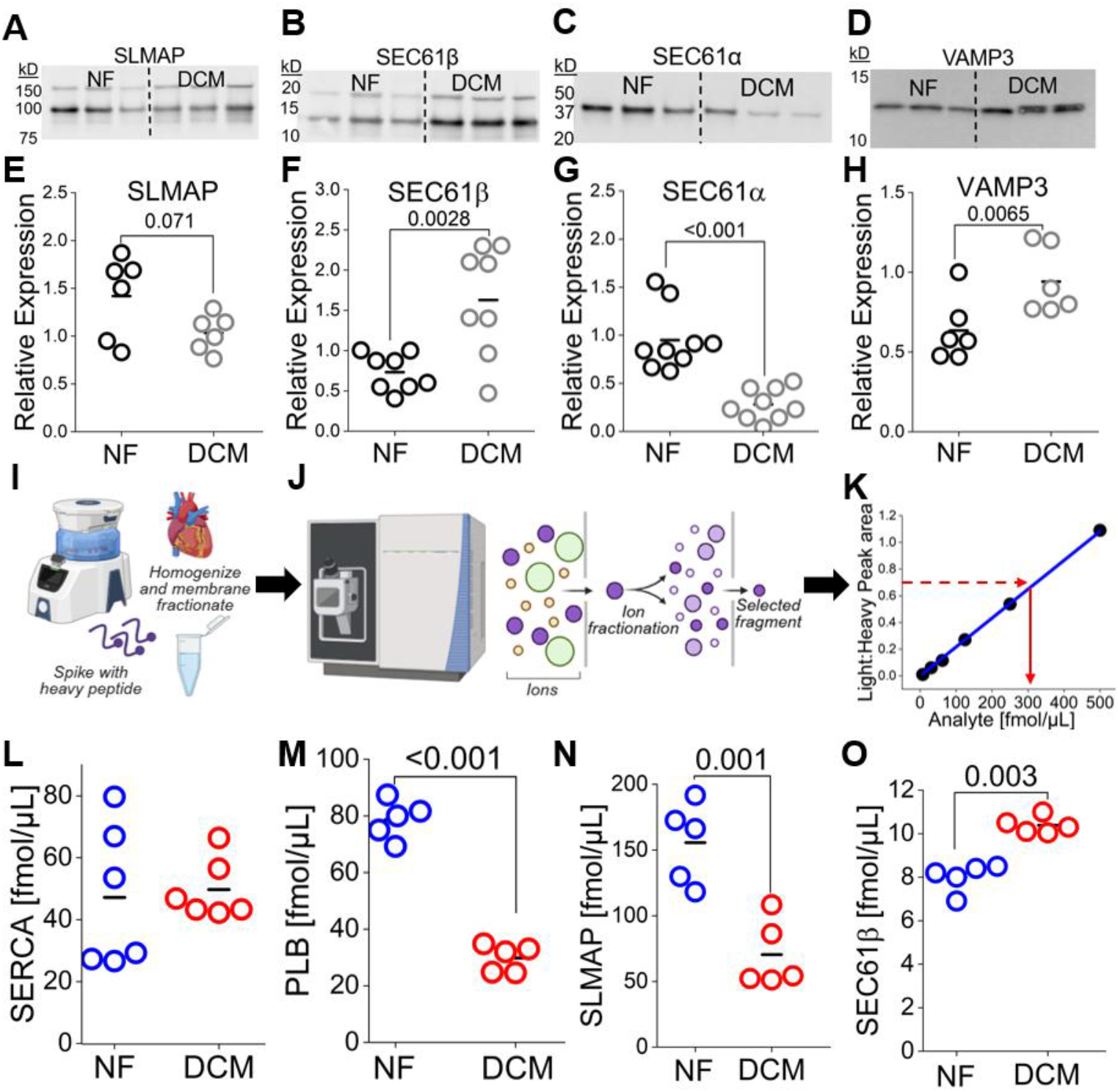
**A)** Representative western blot of SLMAP. **B)** Representative western blot of SEC61β subunit. **C)** Representative western blot of SEC61α subunit. **D)** Representative western blot of VAMP3. **E-H)** Quantification of western blots normalized to total protein stain. **I)** MRMs were performed on homogenized and membrane enriched DCM and NF human samples. Samples were then spiked with heavy peptide for labeling and analyzed via MS. **J)** Schematic for the ion and fragment selection that is required for multiple reaction monitoring (MRM). **K)** A standard curve was created for each protein of interest by varying the ratio of light to heavy peptide and used to interpolate analyte concentration in each sample. **L)** SERCA displays no change in expression between NF and DCM samples and has a concentration of around 50 fmol/µL **M)** PLB decreases from 80 fmol/µL to 30 fmol/µL in DCM. PLB samples were treated with alkaline phosphatase, so is representative of total PLB. **N)** Full-length SLMAP decreases from 150 fmol/µL to 75 fmol/µL in DCM. **O)** SEC61β increases from 8 fmol/µL to 11 fmol/µL in DCM.

While peptide candidates exhibited increased relative expression in heart failure, if their absolute levels are far below that of SERCA, their impact is likely to be modest at best. Thus, to quantify the absolute concentration of the most inhibitory candidates, we performed quantitative mass spectrometry using multiple reaction monitoring (MRM), as described in *Materials and Methods*. Briefly, we synthesized a heavy and light form of each candidate of interest, based on the peptide sequence detected for that candidate in the discovery mass spectrometry experiments. Heavy peptides are composed of the same amino acid sequence as the light peptide but are isotopically distinguishable on the mass spectrometer due to substitution of deuterium for hydrogen on the N-terminal residue. A standard curve was made for each candidate using varying ratios of heavy to light peptides (**Fig. 5K**) for calibration of a triple quadrupole mass spectrometer. A known quantity of heavy peptide was added to specimens (**Fig. 5I**), then exogenous heavy peptide and endogenous light peptide were detected and quantified for each sample to measure the absolute quantity of each candidate poison peptide in the specimens (**Fig. 5J**). Like discovery mass spectrometry experiments, the non-failing and DCM human heart specimens were membrane enriched to increase sensitivity to membrane bound proteins. Unlike discovery mass spectrometry experiments, these MRMs were conducted on tissue lysates not subjected to size restriction. This was done for two primary reasons: MRM assays require a large sample concentration to accurately quantitate abundance and require consistent sample preparation to be compared to the other protein abundances. Therefore, these measurements are reflective of total full-length protein in each sample rather than possible degradation products from smaller gel bands.

MRM analysis revealed that SERCA expression was unchanged in DCM, remaining at an average of 50 fmol/µL across samples (**Fig. 5L**), consistent with recent literature (Ragone et al. 2023). Of note, PLB showed a marked change in DCM compared to non-failing hearts, decreasing from 80 fmol/µL to 30 fmol/µL (**Fig. 5M**). The data are consistent with a decrease in the PLB:SERCA ratio from 1.6 in non-failing hearts to 0.6 in DCM hearts. Full-length SLMAP was shown to have a significant decrease in expression in DCM (**Fig. 5N**), consistent with the degradation of full-length protein and production of smaller peptide fragments we saw in discovery mass spectrometry at smaller gel sizes (**Fig. 1B**). Abundance decreases by around 75 fmol/µL in DCM, suggesting that up to 75 fmol/µL of transmembrane SLMAP fragment could be produced. SEC61β showed a significant increase in DCM to around 11 fmol/µL (**Fig. 5O**). Overall, through MRM we see an increase in these toxic peptides and conclude their absolute abundance is near or above that of SERCA and PLB. This indicates that their impact in vivo is substantial and contributing to the SERCA dysregulation and calcium mishandling observed in DCM.

## Discussion

### Toxic competitors of PLB contribute to dysregulation of SERCA

The present study revealed a novel pathophysiological mechanism that may contribute to calcium mishandling in the heart: toxic peptides that bind to SERCA, displacing endogenous regulatory species and inhibiting the pump without responsiveness to native signaling pathways (**Fig. 6A**). Several pieces of evidence support the hypothesis that this mechanism is operative in the failing heart. In cardiac specimens from patients with DCM we observed striking alterations to transcription and proteostasis, resulting in generation of many small peptides through increased synthesis and by proteolytic cleavage of larger precursors. Of the 92 proteins found to be significantly upregulated in DCM, many contain multiple transmembrane helices that could be considered individual candidates. From the list of peptides we examined here, half bound avidly to the SERCA pump, and many of these showed potent inhibition of SERCA transport activity. This suggests that there may be many more toxic peptides, beyond those screened in this study, contributing to calcium mishandling observed in DCM.

**Figure 6.**
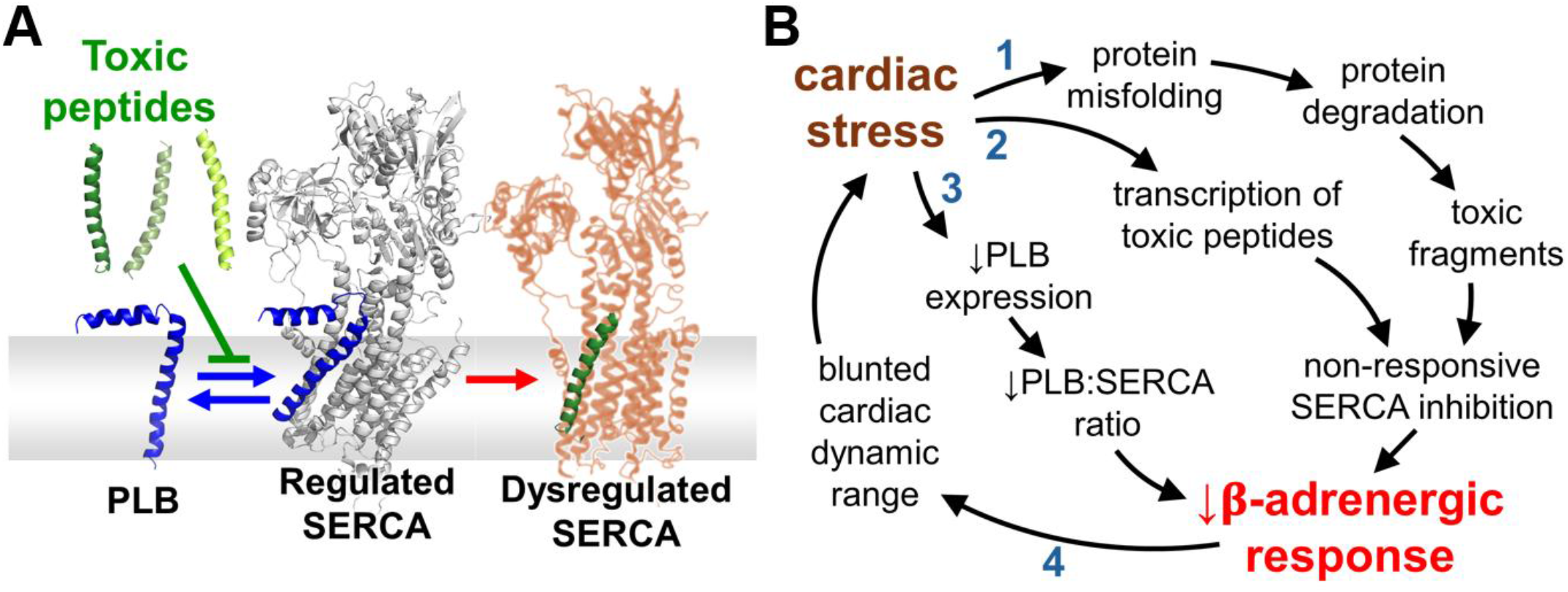
**A)** Toxic peptides directly compete with PLB mediated regulation of SERCA leading to severe dysregulation of the pump. **B)** Proposed mechanism for SERCA dysregulation in DCM due to production of toxic peptides. Loss of SERCA response to β-adrenergic signaling further increases cardiac stress and accelerates disease progression.

### Pathogenic cycle of Ca handling disruption

**Fig. 6B** outlines how this pathological process could contribute to the degradation of cardiac function during the evolution of disease. The process of heart failure can begin from a variety of causes, including diverse genetic lesions^32,46-48^, metabolic dysfunction^49,50^, or mechanical stress from structural disease or hypertension^2,51^. The heart has some capacity to adapt to such stresses, but once cardiac stress becomes maladaptive, the cycle of declining cardiac function begins. Protein misfolding results in increased protein degradation, which soon overwhelms the proteosome, leading to accumulation of toxic protein fragments (**Fig. 6B-1**). Toxic peptides are also produced directly from transcriptional upregulation (**Fig. 6B-2**). Together with proteolytic fragments, they compete with native regulatory micropeptides such as PLB for binding to SERCA. In addition to competitively displacing PLB, some of the candidates demonstrate a direct functional inhibition of SERCA, reducing Ca uptake (**Fig.4A**) and decreasing the rate of maximal turnover (**Fig.4D**). Lacking the PKA and CamKII phosphorylation sites of PLB, toxic peptide species cannot respond to adrenergic signaling. Toxic competition of SERCA-binding peptides for SERCA is exacerbated by a decline in PLB protein expression (**Fig. 6B-3**), exposing a greater population of SERCA to dysregulation. If SERCA becomes inhibited and unresponsive to native regulatory, the heart’s dynamic range will be blunted. Poor regulation of cardiac performance between rest and stress reinforces the maladaptive stress response (**Fig. 6B)**.

### Reciprocal changes in peptide species

One might reasonably expect that, during DCM, downregulation of the native inhibitor PLB might be functionally offset by the increase in other inhibitory peptide species produced by protein cleavage or increased transcription (**Fig. 6**). However, we postulate that toxic peptide inhibition cannot recapitulate native inhibitory mechanisms. As described above, the extraneous peptide species produced in DCM are presumably unresponsive to kinase regulation. Moreover, SERCA regulation by PLB is precisely tuned to respond to other cellular signals. PLB inhibits SERCA by reducing the pump’s Ca^2+^ affinity, without significantly reducing turnover rate^9,52-54^. Thus, SERCA is most strongly inhibited when cytoplasmic Ca^2+^ is low, preventing excessive and unnecessary SERCA cycling at the end of the diastolic (relaxation) phase of the cardiac cycle. Then, PLB inhibition of SERCA is fully relieved when Ca levels rise during the systolic (contraction) phase^55^, allowing SERCA to still achieve its maximal turnover rate in spite of PLB. This important relief-of-inhibition mechanism appears to be absent for at least one of the toxic peptides investigated here, SEC61β. ATPase measurements showed that this toxic peptide does not alter SERCA Ca^2+^ sensitivity; instead, the observed reduction in Ca^2+^ uptake (**Fig. 4A, C**) is due to a decrease in maximal SERCA turnover rate (**Fig. 4D**). These characteristics predict insufficient suppression of SERCA activity at low Ca^2+^, and sluggish SERCA activity at high Ca^2+^, when it is needed most. This inhibitory profile would damp changes in Ca^2+^ handling between diastole and systole or between rest and exercise, thereby contributing to the loss of cardiac dynamic range (**Fig. 6B-4**).

### Relevance to human disease

It is a limitation of the present study that MS analysis was performed on explanted DCM hearts representing end-stage failure, in which many aspects of cardiac function (besides proteostasis) are severely compromised. It is difficult to attribute late-stage dysfunction to any pathophysiological mechanism, and the extent to which toxic peptides may contribute to the evolution of heart failure is unknown. However, it is noteworthy that many of the candidate toxic peptides were also detected in complementary MS analyses of specimens of an animal model of mild heart failure (**Fig.1G**). This TAC model was monitored via echo cardiography and compared to SHAM control, displayed no change to ejection fraction but a moderate increase in left ventricle diameter and wall thickness (**Fig. 1E-F**). We consider this model to be representative of an early stage of heart failure, prior to decompensation. Elevated levels of candidate toxic peptides in this model suggest that this process is initiated early in the evolution of disease, which is compatible with the hypothesis that aberrant SERCA interactions with toxic species contribute to the worsening of cardiac function throughout the disease process.

### Toxic Peptides: A General Pathophysiological Mechanism?

This mechanism may not be unique to the SERCA pump and has the potential to affect many other P-type pumps with broadly specific regulatory sites. Plasma membrane (PM) transporters may also encounter toxic peptides, as we detected many degraded PM-localized peptides by mass spectrometry. We posit that any protein with a broadly specific regulatory site in the transmembrane region may be susceptible to this pathological mechanism. As a likely example, the sodium potassium ATPase (NKA) is also regulated by a wide range of regulatory partners (FXYD proteins)^56^. Although each pump may have their own motif of what is able to occupy the binding site, the diverse alterations to proteostasis and transcriptional regulation that occur in disease, lead to a wide array of small peptide production. This upregulation of small transmembrane peptides may compete with each respective regulatory mechanism and lead to disruption of pump activity and function.

### Summary

We conclude that small inhibitory peptides, produced during periods of cardiac stress, are contributing to the calcium mishandling and loss of the β-adrenergic response that are observed in DCM. We have identified three specific peptides that on their own lead to excessive SERCA inhibition, but posit there are many other candidates whose additive inhibitory effect further contribute to this mechanism. This is a novel mechanism that should be explored for therapeutic intervention to restore proper cardiac relaxation in DCM.

## Materials and Methods

### Discovery mass spectrometry

#### Sample preparation

Left ventricle samples from non-failing and DCM human hearts were flash frozen and stored at -80°C, previously. Non-failing samples were taken from rejected donors with cause of death due to non-cardiovascular related issues and DCM samples were from end stage heart failure patients. Samples were thawed and homogenized (Omni international GLH 850) in SR prep buffer (20 mM MOPS, 0.1 M KCl, PH=7). Samples were centrifuged at 4,100 rcf for 20 minutes at 4°C. Supernatant was collected and pellet was resuspended in SR prep buffer to be homogenized and centrifuged a second time at 4,100 rcf for 30 minutes. Combined supernatants from both centrifugation steps were then spun at 54,000 rcf for 1 hour (Beckman Coulter Optima TLX Ultracentrifuge). Pellet was resuspended in sucrose buffer (20 mM MOPS, 0.2 M Sucrose, pH=7) and homogenized with a Dounce homogenizer to obtain SR membrane fractions. A BCA was performed to determine protein concentration and samples can be stored at -80°C.

10 μg of triplicate samples of non-failing and DCM membrane fractions boiled for 30 minutes at 65°C and mixed with Laemmli buffer. Samples were run on an SDS-PAGE gel (10% acrylamide stacking layer, 20% acrylamide resolving layer) next to a 10kDa protein ladder and 1.5 kDa low MW protein ladder for 16 hours at 200V. The gel was then stained with oriole fluorescent gel stain for 1 hour and imaged under UV light. Sections of gel were cut using razorblades above and below 26kDa and placed in 1.5 mL tubes. Gel pieces were destained with 50% ACN and 50% 50 mM NH4HCO3 followed by 100% ACN and dried on a speedvac. Gel pieces were reduced in DTT, alkylated in IAA, and dehydrated with ACN. Gel pieces were then covered with trypsin (12.5 ng/uL) at 37°C overnight to digest sample. Digested proteins were extracted from gel using buffer A (1% formic acid, 50% ACN) and dried using a speedvac. Pierce peptide quantification assay was used to determine peptide concentration of sample and then 20uL of sample was resuspended in buffer A.

#### Proteomics

Purified peptides, 500 ng, were loaded onto a Vanquish Neo UHPLC system (Thermo Fisher) with a heated trap and elute workflow with a c18 PrepMap, 5mm, 5uM trap column(P/N 160454) in a forward-flush configuration connected to a 25cm Easyspray analytical column(P/N ES802A rev2) 2uM,100A,75um x 25 with 100% Buffer A (0.1% Formic acid in water) with flow rate of 0.300 µL and the column oven operating at 35 °C. Peptides were eluted over a 90 min gradient, using 80% acetonitrile 0.1% formic acid (buffer B), going from 2.5 % to 8% over 5 min, to 30% over 55 min, then to 45% over 15 min, then to 99% over 8 min and kept at 99% for 6 min, after which all peptides were eluted. Spectra were acquired with an Orbitrap Eclipse Tribrid mass spectrometer (Eclipse) with FAIMS Pro interface (Thermo Fisher Scientific) running Tune 3.5 and Xcalibur 4.5 was used for all acquisition methods, spray voltage set to 2000V, and ion transfer tube temperature set at 300oC, FAIMS switched between CVs of −45 V, – 55 V, and −65 V with cycle times of 1.5 s. MS1 spectra were acquired at 120,000 resolutions with a scan range from 375 to 1500 m/z, normalized AGC target of 300%, and maximum injection time set to auto, S-lens RF level set to 30 without source fragmentation and datatype positive and profile; Precursors were filtered using monoisotopic peak determination set to peptide MIPS; included charge states, 2-7 (reject unassigned); dynamic exclusion enabled, with n = 1 for 60s exclusion duration at 10 ppm for high and low. DDMS2 scan using isolation mode Quadrupole, Isolation Window (m/z): 1.6; Activation Type set to HCD with 30% Collision Energy (CE), Detector Type: Ion Trap; Scan Rate: Rapid; AGC Target: 10000; Maximum Injection Time:35 ms, Microscans: 1 and Data Type: Centroid.

#### MS Data Analysis

Mass spectrometry chromatograms were analyzed using Proteome Discoverer 2.5 (Thermo Fisher) using Sequest HT search engines. The data were searched against the Human entries in the UniProt protein sequence database (Homo sapiens, Proteome ID UP000005640). The sequest search parameters included precursor mass tolerance of ten ppm and 0.6 Da for fragments, 2 missed trypsin cleavages, oxidation (Met) and acetylation (protein N-term) as variable modifications, and carbamidomethylation (Cys) as a static modification. Percolator PSM validation was used with the following parameters: strict false discover rate (FDR) of 0.01, relaxed FDR of 0.05, maximum ΔCn of 0.05, and validation based on q-value. Precursor Ions Quantifier settings were Peptides to Use: Unique + Razor; Consider Protein Groups for Peptide Uniqueness set as True; Precursor Abundance Based On: Intensity; Normalization based on Total Peptide Amount; Scaling Mode set as none, Protein Abundance Calculation based on Top 6 Average intensity, low abundance peptides were removed by filtering out proteins with less than 3 PSMs, Pairwise ratio based calculation and T-test(background based) were used for class comparison. Differentially expressed proteins were selected based on p-value < 0.05 and log2 fold change > 1.0.

### Cell culture and transfection

pEGFP-C1 was used as an expression vector and human sequences of all peptide candidates and controls were labeled using YFP on the N-terminal end with a 5 amino acid linker of sequence SGLRS. Peptide sequences were inserted into the vector using Gibson Assembly (New England Biolabs, Ipswich, MA) and custom oligonucleotide primers (Eurofins MWG Operon). SERCA was labeled with mCer on the N-terminal end with the same linker sequence. HEK293Tcells (Agilent, Santa Clara, CA, USA) were cultured in DMEM cell culture medium with 10% fetal bovine serum (Thermo Fisher Scientific, Waltham, MA, USA). The cells were then co-transfected with 0.5 μg mCer-SERCA and 1.5 μg YFP-candidate peptides using Lipofectamine 3000 transfection kit (Invitrogen Life Technologies, Carlsbad, CA, USA) per the manufacturer’s instructions and plated on 60mm cell culture plates. After 48 h of protein expression, the cells were trypsinized (Thermo Fisher Scientific) and replated onto poly-D-lysine-coated 2-well glass bottom chamber plates before imaging. Prior to imaging, cells were incubated at 37C in DMEM with 10% fetal bovine serum.

### Structured illumination microscopy

HEK293T cells were transfected with 1 μg mCer-SERCA and 1 μg YFP-candidates or controls as previously described. After 48 hours of protein expression cells were trypsinized and transferred to a 4 well glass bottom chamber slides. Cells were washed once with PBS prior to imaging. Cells were imaged using a 64X oil immersion objective (1.4 NA) on the Lattice SIM 5 microscope (Zeiss). Images were acquired under 405 nm and 488 nm excitation wavelengths for visualization of mCer and YFP using a Hamamatsu Camera and 50 ms exposure time. 27 μm apotome grating was used and a 20-image z-stack was acquired for each cell. Images were SIM processed using ZEN 3.0 software (Zeiss) set to weak sharpness and fast fit histogram.

### FRET microscopy

Imaging was performed using a wide-field fluorescent microscope as described previously. Cells were imaged in PBS using an inverted microscope (Nikon Eclipse Ti2) equipped with a photometrics camera (PRIME 95B, 25 mm). The image acquisition of a field of view for each sample was performed with a 40 × 0.95 NA objective and 100 ms exposure time for Cer, YFP, and FRET (Cer excitation, YFP emission) channels. Metamorph software (Molecular Devices, Sunnyvale, CA) was used to quantify fluorescent intensity for the images collected in each channel. FRET efficiency was calculated according to *E = G/(G + 3*.*2 × F*_*Cer*_), where *G* = *F*_FRET_ − *a × F*_YFP_ − *d × F*_Cer_, where *F*_FRET_, *F*_YFP_, and *F*_Cer_ are the matching fluorescence intensity from FRET, YFP, and Cer images, respectively, and *G* corrects FRET for crosstalk between channels. The parameters *a* and *d* are crosstalk constants calculated as *a* = *F*_FRET_/*F*_Cer_ for a control sample transfected with only YFP and *d* = *F*_FRET_/*F*_YFP_ for a control sample transfected with only Cer. These values were determined to be *G* = 3.58 *a* = 0.13 and *d* = 0.44. The FRET efficiency of each cell was plotted as a function of protein expression (measured from YFP fluorescence intensity).

### Calcium activity assay

GFP-rCepia1*er* was cloned into DNA plasmids as previous described^43^. HEK293T cells were transfected with 0.5 μg GFP-rCepia1*er*, 2 μg unlabeled RyR2, 0.5 μg unlabeled SERCA2a and 0.5 μg mCer-candidate peptides using lipofectamine 3000 (Invitrogen). Cells were cultured in 60 mm dishes with DMEM and 10% FBS for 48 hours. Cells were incubated in trypsin and counted. Around 100,000 cells were transferred to a 2-well glass bottom imaging dish coated with Poly-D-Lysine and incubated for another hour prior to imaging. Cells were washed once with PBS and then incubated in Tyrode’s buffer (in mM): 140 NaCl, 4 KCl, 2 CaCl2, 1 MgCl2, 10 glucose, and 10 HEPES; pH 7.4 for 10 min prior to imaging to ensure Ca^2+^ equilibration. Each well was scanned and imaged through the 40 × 0.75 numerical aperture objective of a Nikon Eclipse Ti2. For epifluorescence imaging, illumination was introduced through an excitation filter wheel equipped with narrow band filters and a multiple band dichroic mirror. Cells were imaged under 100 ms exposure for each channel: YFP (to image GFP fluorescence) excited using a bandpass excitation filter of 510(25) nm with an emission bandpass filter set of 540(21) nm and mCherry excited using a bandpass excitation filter of 575(25) nm with an emission bandpass filter set of 632(60) nm. Image analysis was performed using FIJI software using a custom macro that selected a mask around GFP expressing cells with an area of 55–2200 μm2 and were at least 40 % circular. GFP and RCEPIA1er fluorescence intensity were then calculated after manual background subtraction and converted to a previously calibrated ER calcium concentration^43^.

### ATPase activity assays

#### Peptide synthesis and purification

Peptides were ordered from and synthesized by Peptide 2.0 (Chantilly, VA). The synthesized peptide sequences are as follows:

Sec61β: KVGPVPVLVMSLLFIASVFMLHIWGK

SLMAP: WMPMLAALVAVTAIVLYVPGL

VAMP3: KMWAIGITVLVIFIIIIIVWVVSS

All three peptides were synthesized with at least 98% purity. Peptides arrived as a lyophilized powder and were resolubilized in 80% isopropyl alcohol, aliquoted (100 μg), dried under vacuum, and stored at -80°C. Recombinant wild-type sarcolipin (wtSLN) was expressed and purified as previously described ^57,58^.

#### Reconstitution of SERCA and peptides into proteoliposomes

For co-reconstitution into proteoliposomes with SERCA, the peptide was resolubilized in 100uL of 80% trifluoroethanol with 375 µg of EYPC and 50 µg EYPA from stock chloroform solutions. Thin films were made by drying the mixture under a gentle stream of nitrogen gas with vortexing and leaving the films under vacuum overnight. Peptide and lipid thin films were hydrated with 94 µL of water and heated (50ºC) for 30 minutes with intermittent vortexing. After cooling to room temperature, C_12_E_8_ was added to a final concentration of 1 mg/mL and vortexed on high for 3 minutes. After vortexing, buffer was added for a final concentration of 20 mM imidazole pH 7, 100 mM KCl, and 0.02% NaN_3_. Detergent-solubilized SERCA1a from rabbit hind leg was purified as previously described ^59^ and added to a concentration of 0.75 mg/mL and equilibrated at room temperature for 15 minutes with gentle stirring. Detergent was slowly removed with the incremental addition of 19 mg of wet SM-2 Biobeads (Bio-Rad, Hercules, CA) over the course of 4 hours. A 20% and 50% sucrose step gradient was used to collect the reconstituted proteoliposomes, with centrifugation for 1 hour at 40 000 x *g*. At the interface of the 20% and 50% layers, a 600 µL aliquot of proteoliposomes was collected, flash frozen, and stored at -80°C until further use.

The co-reconstituted proteoliposomes used in this study approximate the SR membrane in protein density, orientation, and lipid composition ^45,58,60-62^. The proteoliposomes are unilamellar, the SERCA molecules are oriented in the membrane with the cytoplasmic domains accessible to substrates on the outer surface ^60^, and the lipid-to-protein ratio is ∼120 to 1 ^45^. These proteoliposome characteristics mimic native skeletal and cardiac muscle SR membranes, and they allow precise measurement of calcium-dependent ATPase activity (e.g. ^58,61,62^) and calcium transport ^63^.

#### ATPase activity assays

The effect of the peptides and wtSLN on the ATPase activity of SERCA was measured using a coupled-enzyme assay ^64^. The assay was adapted to a 96-well plate format using a Biotek Epoch 2 microplate reader (Agilent, Santa Clara CA). The buffer composition of each well was 50 mM imidazole pH 7, 100 mM KCl, 5 mM MgCl_2_, 2.4 mM ATP, 0.5 mM EGTA, 0.5 mM PEP, and 0.18 mM NADH in a 160 µL well volume. Each well contained 9.6 units/mL of lactate dehydrogenase and pyruvate kinase. The calcium-dependent ATPase activity of SERCA was measured by the conversion of NADH to NAD^+^ coupled to ATP hydrolysis, monitored by measuring the absorbance at 340 nm. All assays were done with proteoliposomes containing SERCA in the absence and presence of the poison peptides or wtSLN. The proteoliposomes were diluted with 20% w/v sucrose (20 mM imidazole pH 7, 100 mM KCl, 0.02% NaN_2_) at a 1:5 volume ratio and 3.2 nM calcium ionophore was added. The 96-well plates comprised a range of calcium (0.1 to 10 µM) and SERCA (10 to 20 nM) concentrations. Assays were conducted at 30°C, initiated *via* the addition of proteoliposomes to the wells, and data points at 340 nm were collected every ∼50s for 30 minutes. These plots were linear for the time frame of the experiments. At least three independent reconstitutions and assays were done on each set of proteoliposomes. Sigma Plot software (SPSS Inc., Chicago, IL) was used to calculate the V_max_, K_Ca_, and n_H_ *via* the non-linear least-squares fitting of the Hill equation to the activity data. Errors were calculated as the standard error of the mean. Comparison of the K_Ca_ and V_max_ parameters was carried out using between-subjects, one-way analysis of variance followed by the Holm-Sidak test for pairwise comparisons.

### Multiple reaction monitoring (MRM)

#### Standard curve

Heavy and light peptides were synthesized of each protein of interest (AQUA thermofisher). Peptides were direct injected into the Altis mass spectrometer to determine product ions, collision energy and RF lens optimal settings for each set of peptides. Custom injection methods were created for each compound. Varying ratios of heavy and light peptide were then run on the Atlis to create a standard curve (**Fig. 5K**) The amount of heavy peptide was kept constant at 500 fmol while serial dilations of light peptide were used from 7.5 to 500 fmol/uL.

#### Sample Preparation

SR membrane fractions from non-failing and DCM human heart samples were prepared as previously described. A chloroform methanol precipitation was used to remove sucrose from sample and DTT and IAA were used for reduction and alkylation, respectively. Final concentration of 500 fmol/μL of heavy peptide was spiked into each sample. 1 μg of trypsin in Tris-HCl was added and samples were incubated overnight at 37°C. A Pierce desalting column (protocol included in kit) was used to collect peptide fragments; samples were dried with a speed vac and resuspended in 40 μL buffer A. A pierce peptide quantification assay was performed to determine concentration.

#### Proteomics

Purified peptides, 500 ng, were loaded onto a Dionex UltiMate 3000 RSLCnano HPLC System (Thermo Fisher) with a heated trap and elute workflow with a c18 Prep Map, 5mm, 5uM trap column(P/N 160454) in a forward-flush configuration connected to a 25cm Easy spray analytical column(P/N ES802A rev2) 2uM,100A,75um x 25 with Buffer A with flow rate of 0.350 µL and the column oven operating at 35 °C. Peptides were eluted over a 60 min gradient, using 80% acetonitrile 0.1% formic acid (buffer B), going from 5 % to 10% over 8 min, to 45% at 45 min, then to 98% at 49 min, and kept at 98% for 11 min, after which all peptides were eluted. Spectra were acquired with TSQ Altis, Triple Quadrupole MS instrument running Tune 3.5 and Xcalibur 4.5 were used for all acquisition methods, spray voltage set to 1700V, and ion transfer tube temperature set at 300°C. Collision energy (V) and RF lens (V) varied between peptides and are listed in the table below:

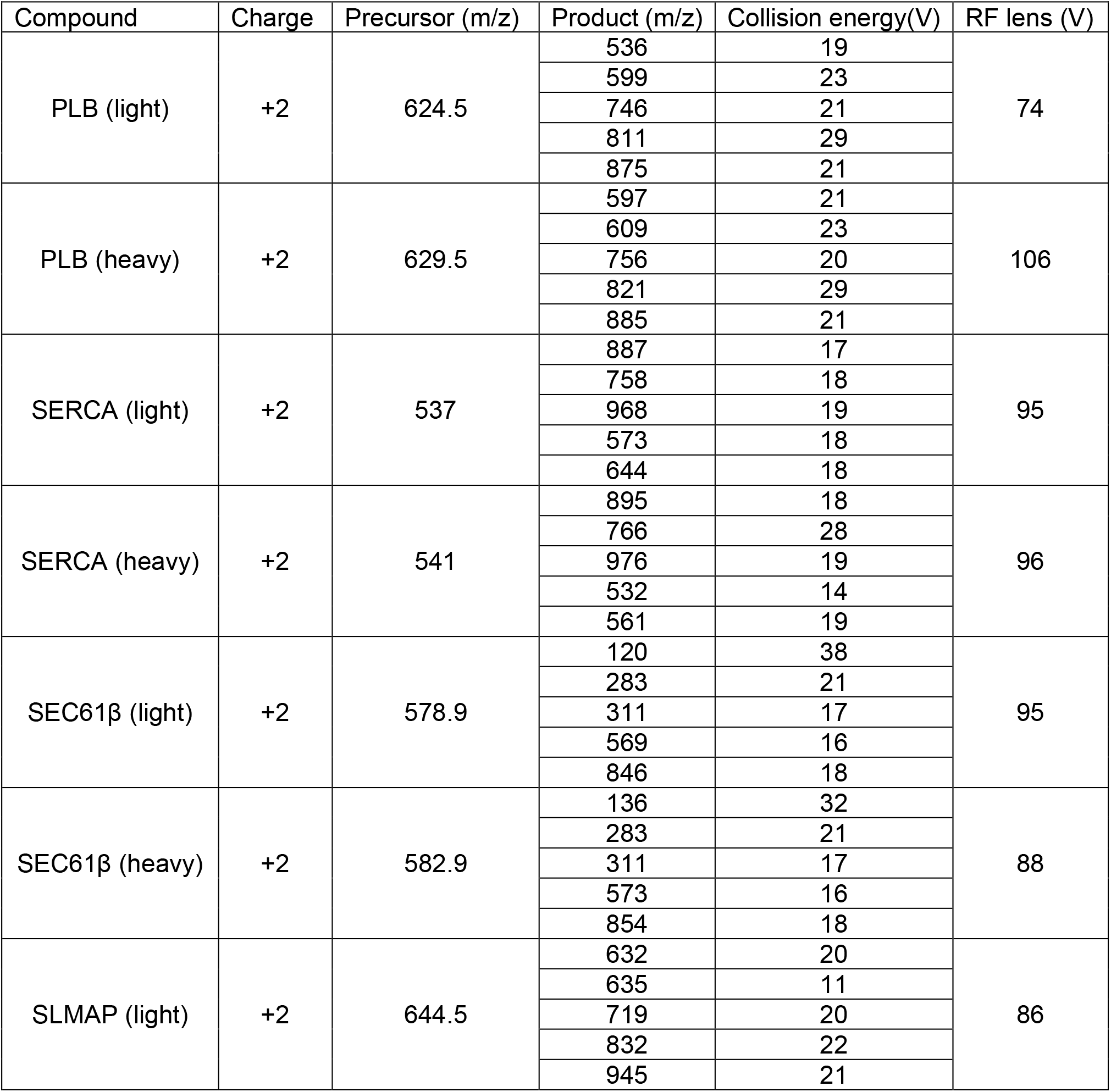
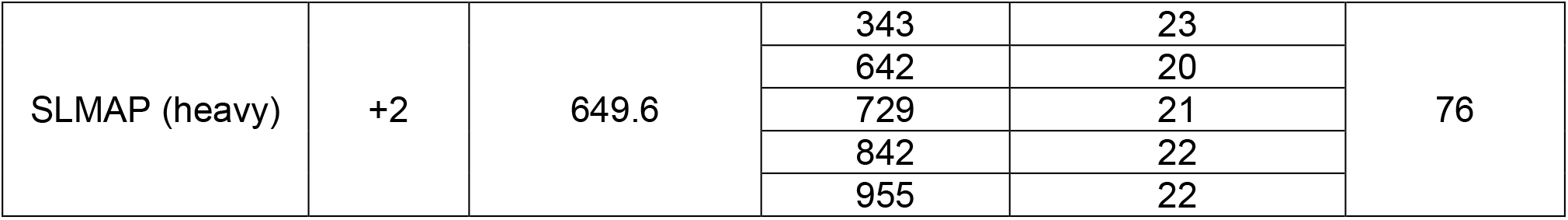

| Compound | Charge | Precursor (m/z) | Product (m/z) | Collision energy(V) | RF lens (V) |
| --- | --- | --- | --- | --- | --- |
| PLB (light) | +2 | 624.5 | 536 | 19 | 74 |
|  |  |  | 599 | 23 |  |
|  |  |  | 746 | 21 |  |
|  |  |  | 811 | 29 |  |
|  |  |  | 875 | 21 |  |
| PLB (heavy) | +2 | 629.5 | 597 | 21 | 106 |
|  |  |  | 609 | 23 |  |
|  |  |  | 756 | 20 |  |
|  |  |  | 821 | 29 |  |
|  |  |  | 885 | 21 |  |
| SERCA (light) | +2 | 537 | 887 | 17 | 95 |
|  |  |  | 758 | 18 |  |
|  |  |  | 968 | 19 |  |
|  |  |  | 573 | 18 |  |
|  |  |  | 644 | 18 |  |
| SERCA (heavy) | +2 | 541 | 895 | 18 | 96 |
|  |  |  | 766 | 28 |  |
|  |  |  | 976 | 19 |  |
|  |  |  | 532 | 14 |  |
|  |  |  | 561 | 19 |  |
| SEC61β (light) | +2 | 578.9 | 120 | 38 | 95 |
|  |  |  | 283 | 21 |  |
|  |  |  | 311 | 17 |  |
|  |  |  | 569 | 16 |  |
|  |  |  | 846 | 18 |  |
| SEC61β (heavy) | +2 | 582.9 | 136 | 32 | 88 |
|  |  |  | 283 | 21 |  |
|  |  |  | 311 | 17 |  |
|  |  |  | 573 | 16 |  |
|  |  |  | 854 | 18 |  |
| SLMAP (light) | +2 | 644.5 | 632 | 20 | 86 |
|  |  |  | 635 | 11 |  |
|  |  |  | 719 | 20 |  |
|  |  |  | 832 | 22 |  |
|  |  |  | 945 | 21 |  |
| SLMAP (heavy) | +2 | 649.6 | 343 | 23 | 76 |
|  |  |  | 642 | 20 |  |
|  |  |  | 729 | 21 |  |
|  |  |  | 842 | 22 |  |
|  |  |  | 955 | 22 |  |

#### MS Analysis

Elution time and absolute abundance, measured by area under the curve for each peak, was quantified using Skyline (Thermo Fisher) MS Software. A standard curve was created for each protein with heavy peptide concentration kept constant at 500 fmol/μL and light peptide ranging from 7.5-500 fmol/μL. Abundance of light peptide product present in each sample was plotted on the standard curve and final concentration was calculated (N=6 non-failing and DCM samples used for each MRM)

### Statistical Analysis

Power analysis was performed to calculate sample size for each experiment. Errors are reported as standard errors, and statistical significance was evaluated using one-way analysis of variance (between subjects and samples) followed by Tukey’s post hoc where p < 0.05 was considered significant using the software ORIGIN (OriginLab, Northampton, MA).

## Notes

### Competing Interest Statement

The authors have declared no competing interest.

### Funding

- National Heart Lung and Blood Institute: HL092321, HL158649, HL143816

